# Chronic consumption of alcohol increase alveolar bone loss

**DOI:** 10.1101/2020.04.22.055186

**Authors:** Juliano Milanezi de Almeida, Victor Fabrizio Cabrera Pazmino, Vivian Cristina Noronha Novaes, Suely Regina Mogami Bomfim, Maria José Hitomi Nagata, Fred Lucas Pinto Oliveira, Edilson Ervolino

## Abstract

**Background:** This study evaluated the effects of the chronic consumption of different concentrations of alcohol on the experimental periodontitis (EP).

**Methods:** 160 rats were divided into 4 groups: (EP-NT) rats with EP and no alcohol exposure; (EP-A14) rats with EP exposed to 14% alcohol; (EP-A25) rats with EP exposed to 25% alcohol; (EP-A36) rats with EP exposed to 36% alcohol. The animals from the EP-A14, EP-A25 and EP-A36 groups were subjected to different concentrations of alcohol 30 days before EP induction. The histological characteristics, percentage of bone in the furcation (PBF) and bone metabolism in the furcation region were evaluated. The PBF and tartrate-resistant acid phosphatase (TRAP) data were subjected to statistical analysis.

**Results:** The EP-A14, EP-A25 and EP-A36 groups had lower PBFs compared with the EP-NT group. A more severe inflammatory process and a greater number of TRAP+ cells were also observed. In the EP-A14, EP-A25 and EP-A36 groups, the inflammatory process became more severe as the ingested alcoholic concentration increased. An increase in RANKL immunostaining and a significantly higher number of TRAP+ cells were also observed.

**Conclusion:** We conclude that chronic alcohol consumption increases the severity of experimental periodontitis in a dose-dependent manner by increasing the magnitude of local inflammatory responses and stimulating alveolar bone resorption.

## 1. Introduction

Periodontitis is a multifactorial infectious-inflammatory disease [1], and its progression may be affected by local and systemic risk factors, including social and behavioral factors such as smoking and alcohol abuse [2,3]. The assessment of risk factors associated with periodontitis is of great importance when determining treatment and prognosis. Many clinical studies have evaluated the relationship between alcohol abuse and the risk of periodontal disease and have reported some evidence of an association [4–9]. Based on their meta-analysis from Wang et al., (2016) [10] concluded that alcohol consumption was associated with an increased risk of periodontal disease and must be considered a behavioral risk factor.

The consumption of alcohol is a socially accepted habit and is part of most people’s lifestyle. However, alcohol abuse is considered by the World Health Organization (WHO, 2014) [11] to be the third leading cause of death worldwide, preceded only by cancer and cardiovascular disease. Alcohol abuse has negative effects on systemic health, with liver cirrhosis being the most common chronic disease resulting from alcohol hepatotoxicity [12]. In addition, alcohol directly affects the immune system, increasing the risk of severe infections and altering the metabolic functions of bone homeostasis. These effects, together with the pathology of periodontitis, can accelerate the alveolar bone resorption process, thereby affecting the inflammatory response mechanism of periodontitis [13–17].

The deleterious effects of chronic alcohol consumption have been suggested to be dose dependent [6,8]. In a clinical study, Tezal et al., (2004) [6] observed a significant relationship between the number of drinks per week (5, 10, 15 or 20) and clinical attachment loss (CAL). Lages et al., (2012) [8] observed a higher incidence of periodontitis that corresponded to the frequency of alcohol consumption. According to their results, the prevalence of periodontitis was 53% in alcohol-dependent individuals, and prevalence of 17.2%, 24.0% and 29.6% were observed in occasional, moderate and intense alcohol users, respectively. In a recent study with data from the National Health and Nutrition Examination Survey (NHANES, 2009–2012), Gay et al., (2018) [18] concluded that alcohol consumption was associated with an increased chance of having periodontitis. However, consumption of <1 drink per week was associated with similar odds of having periodontitis compared with non-alcohol consumption. However, the exact dose required to affect the pathology of periodontitis remains unclear.

The mechanisms by which chronic alcohol consumption affects the progression of periodontal disease require further investigation. With respect to bone tissue, studies have shown that chronic alcohol consumption can affect bone density, calcium and phosphorus levels [19,20] and inhibit bone repair [21]. Dal Fabbro et al., (2019) [22], in a recent study concluded that chronic alcohol consumption had a significant effect on the severity of apical periodontitis, exacerbating the inflammatory response and osteoclastogenesis. Consequently, such events resulting from chronic alcohol consumption can compromise the stability of periodontal tissue [6,8,23–26]. Moreover, in a study using animal models, Liberman et al., (2011) [27] concluded that ingestion of alcohol at low concentrations (5%) did not affect alveolar bone loss in induced periodontal disease, suggesting that the adverse effects of alcohol are related to its concentration. For this reason, further experimental studies are needed to clarify the way in which chronic alcohol consumption can alter the pathogenesis of periodontal disease.

No information is available on the association between chronic alcohol consumption and its dose-dependent effects on the progression of periodontitis in rats, so the aim of this study was to evaluate the effects of chronic alcohol consumption at different concentrations on the progression of experimental periodontitis (EP). Thus, the hypothesis tested in this study was that chronic alcohol consumption would increase the severity of EP regardless of the concentration consumed.

## 2. Materials and Methods

### 2.1. Animals

A total of 160 male Wistar rats weighing between 250 g and 300 g were used. The experimental protocol was approved by the Ethics Committee on Animal Use (Protocol n° 00636-2013). This study was conducted in accordance with ARRIVE (Animal Research: Reporting of *In Vivo* Experiments) [28].

According to a table generated by a computer program, the animals were divided into the following groups: an EP-NT group (n = 40), normal rats with EP; an EP-A14 group (n = 40), rats with EP that were exposed to alcohol at a concentration of 14% (v/v); an EP-A25 group (n = 40), rats with EP that were exposed to alcohol at a concentration of 25% (v/v); and an EP-A36 group (n = 40), rats with EP that were exposed to alcohol at a concentration of 36% (v/v).

### 2.2. Administration of alcohol and blood sample collection

The administration of the alcohol solution was initiated 30 days before the induction of EP, and the animals of the EP-A14°, EP-A25° and EP-A36° groups received 14%, 25% and 36% v/v alcohol solutions, respectively, following the method of D’Souza El-Guindy et al., (2010) [29]. The alcohol administration was continued until the end of the experiment. Blood samples were collected for analysis of the liver enzymes alanine aminotransferase (ALT) and aspartate aminotransferase (AST). The blood samples (2.0 mL) were obtained via cardiac puncture 30 days before ligature placement, on the day of the ligature placement and at the time of euthanasia.

### 2.3. EP induction method

To induce EP, the rats were anesthetized with a combination of 70 mg/kg of ketamine hydrochloride (Vetaset, Zoetis Iowa, USA) and 6 mg/kg of xylazine hydrochloride (Coopazine, Coopers São Paulo, Brazil). Then, #24 cotton ligatures (Corrente Algodão n° 24, Coats Corrente, SP, Brazil) were wrapped around the first lower left molars and secured at the gingival sulcus with surgical knots until the end of the experiment [30].

### 2.4. Histological processing and immunohistochemistry

The specimens were demineralized in 10% ethylenediaminetetraacetic acid (EDTA) and processed in a conventional manner. Semi-serial sections (4 μm) were obtained in the mesiodistal direction, and 5 equidistant sections of each specimen were stained with hematoxylin and eosin (H&E) for histological and histometric analyses. Other sections were subjected to indirect immunoperoxidase staining with the following primary antibodies: anti-tartrate-resistant acid phosphatase (anti-TRAP).(SC-30833, Santa Cruz Biotechnology, Santa Cruz, CA, USA), anti-receptor activator of nuclear factor kappa-B ligand (anti-RANKL) (SC-7628, Santa Cruz Biotechnology, Santa Cruz, CA, USA) and anti-osteoprotegerin (anti-OPG) (SC-8468, Santa Cruz Biotechnology, Santa Cruz, CA, US). The immunohistochemical staining followed the protocol described by Garcia et al., (2013) [31].

### 2.5. Analysis of results

A calibrated examiner who was blinded to the treatments performed all of the analyses.

### 2.6. Analysis of ALT and AST

Serum ALT and AST levels were assessed to evaluate liver abnormalities resulting from chronic alcohol consumption using a kinetic method according to the manufacturer’s recommendations (Labtest®, Lagoa Santa-MG, Brazil). The samples were analyzed using a LabQuest semi-automatic biochemistry analyzer (Bioplus, Barueri-SP, Brazil).

### 2.7. Histological analysis

A certified histologist (EE) evaluated the following histological parameters according to the protocol proposed by Almeida et al., (2015) [32]: 1) the nature and degree of inflammation; 2) the extent of the inflammatory process; 3) the presence and extent of tissue necrosis; 4) the presence, extent and nature of bone, cementum and dentin resorption; 5) the status of the vasculature; 6) the structural pattern of the extracellular matrix of the periodontal tissue; and 7) the cellularity pattern of the periodontal tissue.

### 2.8. Histometric analysis of percentage bone in the furcation region (PBF)

Another examiner (VFCP) performed the histometric analysis. An image analysis system (Axiovision 4.8.2, Carl Zeiss MicroImaging GmbH, 07740 Jena, Germany) was used to measure (in square millimeters) the total furcation area, the bone area in the furcation region and the PBF (calculated according to the method of Garcia et al. (2015) [33].

### 2.9. Immunohistochemical analyses

A certified histologist (EE) performed the immunohistochemical analyses. A semi-quantitative analysis of RANKL and OPG immunoreactivity pattern in the furcation area was performed with 400x magnification. Three histological sections were used for each animal, and the following immunostaining criteria were used based on the protocol described by Garcia et al., (2015) [33]: 0, total absence of immunoreactivity pattern (IR) cells; 1, low immunoreactivity pattern (approximately 25% IR cells); 2, moderate immunoreactivity pattern (approximately 50% IR cells); and 3, high immunoreactivity pattern (approximately 75% IR cells). A quantitative analysis was performed for TRAP using 5 sections from each animal. The TRAP+ multinucleated cells per square millimeter were counted in a 1000 x 1000 μm area at the center of the interradicular septum at 200x magnification. The boundary of this area was the outer contour of the alveolar bone tissue in the furcation region extending apically to 1000 μm.

### 2.10. Examiner calibration

All the above-identified examiners were blinded to the experimental groups prior to histometric and immunohistochemical analyses. The examiners were trained, and the PBF and TRAP+ cell measurements were carried out in duplicate. The paired t-test was used to calculate the intraexaminer error. A P value > 0.05 in the paired t-test was used to estimate the feasibility of the proposed method.

### 2.11. Statistical analysis

With a sample size of 10 (P = 0.05), the study power was 89%. The collected data were analyzed using software (BioStat version 5.0, Belém, PA, Brazil). The normality of the histometric and immunohistochemical data was verified using the Shapiro-Wilk test (P ≤ 0.05). When a significant difference was indicated by analysis of variance (ANOVA), multiple comparisons were performed using the Tukey test (P ≤ 0.05).

## 3. Results

### 3.1. Analysis of ALT and AST

The ALT results are shown in Figure 1A. In the intragroup analysis, the EP-A14, EP-A25, EP-A36 groups showed higher levels of ALT on days 0, 3, 7, 15 and 30 compared to baseline (p≤0.05). EP-A14, EP-A25 groups showed higher levels of ALT on days 7, 15 and 30 compared to 0 day (p≤0.05). EP-A36 group showed higher levels of ALT on days 3, 7, 15 and 30 compared to 0 day (p≤0.05).

**Figure 1:**
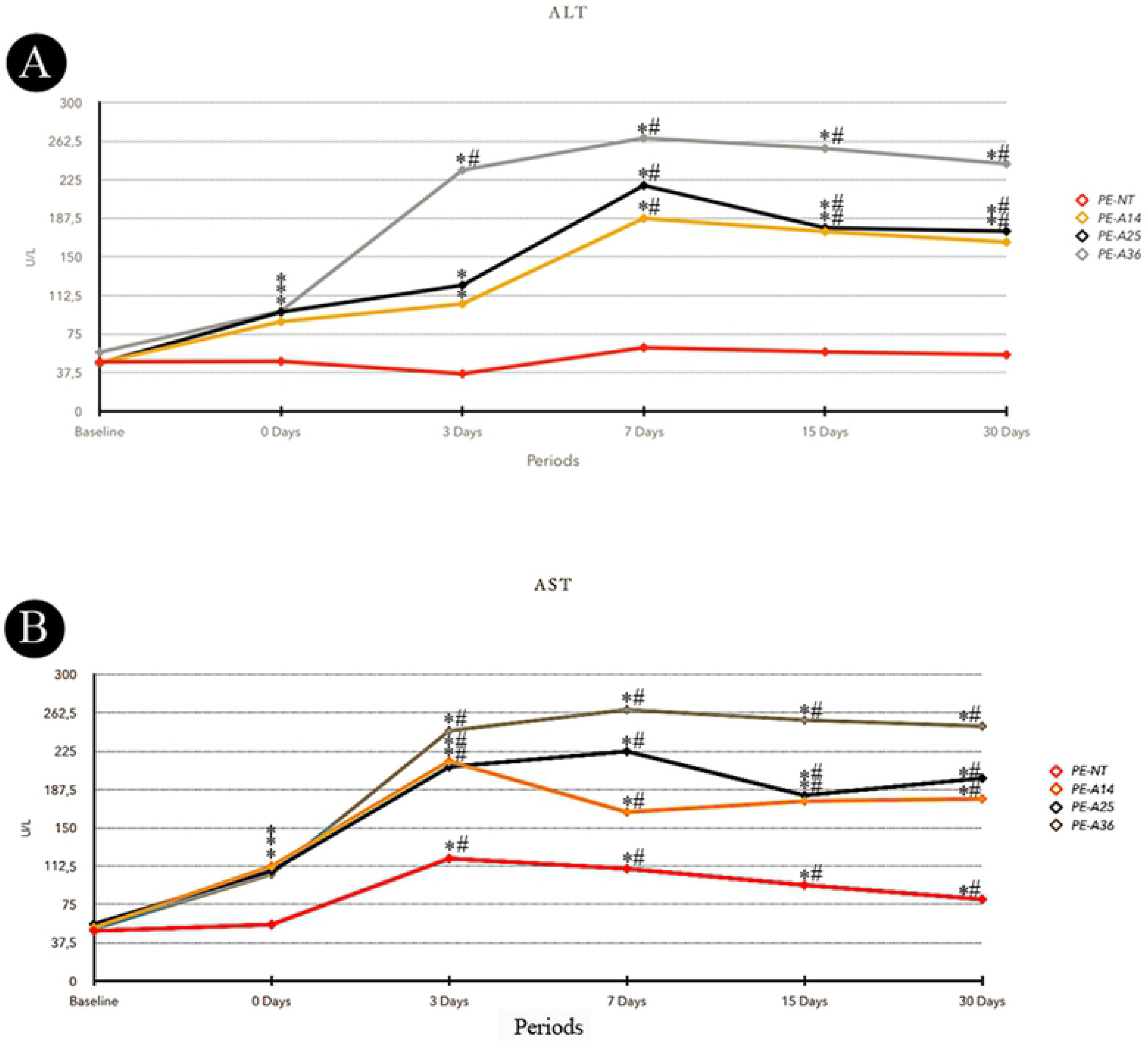
A-Means of ALT serum levels for each group and period. B-Means of AST serum levels for each group and period. Symbols: *statistically significant difference with base line period at the same group. #statistically significant difference with 0 day period at the same group. ANOVA and Tukey test (p≤0.05).

The AST results are shown in Figure 1B. In the intragroup analysis, the EP-NT group showed significant differences, with higher AST levels on days 3, 7, 15 and 30 compared to baseline and day 0 in the same experimental group (p≤0.05). The EP-A14, EP-A25 and EP-A36 groups showed higher levels of AST at day 0 compared to baseline (p≤0.05). At days 3, 7, 15 and 30, the EP-A14, EP-A25 and EP-A36 groups showed higher levels of AST compared to baseline and day 0 of their respective experimental groups (p≤0.05).

### 3.2. Histological analysis

The histological analysis results are shown in Figure 2. Three days after ligature placement, all the experimental groups exhibited severe disorganization of the periodontal ligament in the furcation region. An intense inflammatory infiltrate composed mainly of polymorphonuclear neutrophils was present in this region, and the magnitude of the local inflammatory response was much higher in the groups treated with alcohol. At this time point, a great number of osteoclasts could be observed that were distributed in the furcation region of the alveolar bone, especially in the alveolar crest, which had active bone resorption areas. The levels of bone resorption were similar in all of the experimental groups. However, osteoclast recruitment to the region was significantly increased in the groups treated with alcohol.

**Figure 2:**
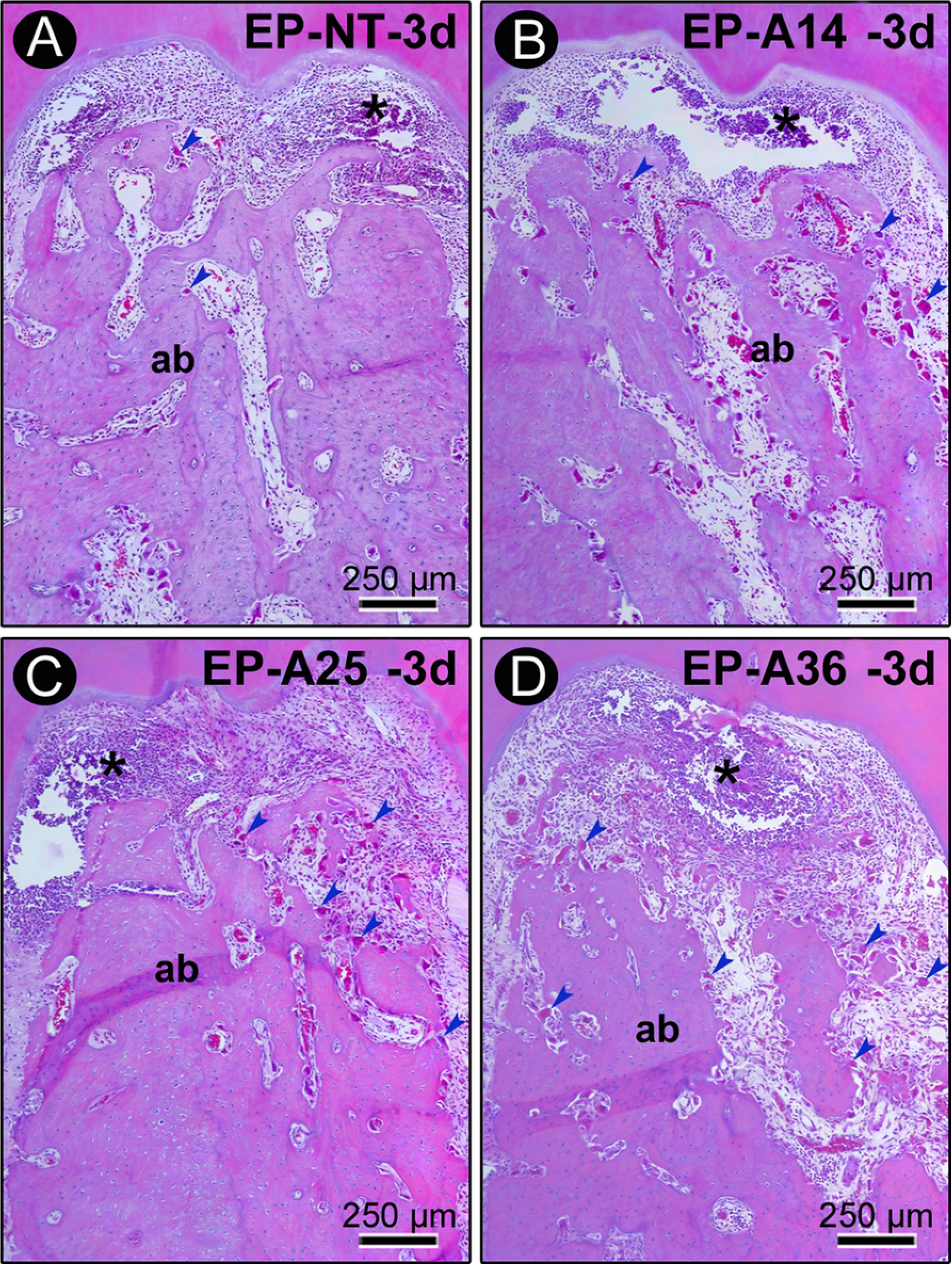
Histopathological characteristics of the furcation region of the mandibular first molars of rats with PE in groups EP (A), EP-A14 (B), EP-A25 (C) and EP-A36 (D) 3 days after ligature placement. Photomicrographs showing the inflammatory infiltrate. Abbreviations and symbols: ab, alveolar bone; *, inflammatory infiltrate. Staining: hematoxylin and eosin (HE). Magnification: 100x. Scale bar: A – d: 250 μm.

Further results are shown in Figure 3. Seven days after ligature placement, tissue disorganization was prevalent in the furcation region, with significantly compromised connective tissue and alveolar bone tissue. Inflammatory infiltration was more extensive, and this inflammatory response was much more exaggerated in the groups treated with alcohol, especially in the EP-A25 and EP-A36 groups. Active bone resorption was present in all specimens from all experimental groups. During this period, the groups treated with alcohol had interradicular septa that consisted of very irregular, extremely thin bone trabeculae with large medullary spaces. In groups EP-A25 and EP-A36, these medullary spaces contained large concentrations of inflammatory cells.

**Figure 3:**
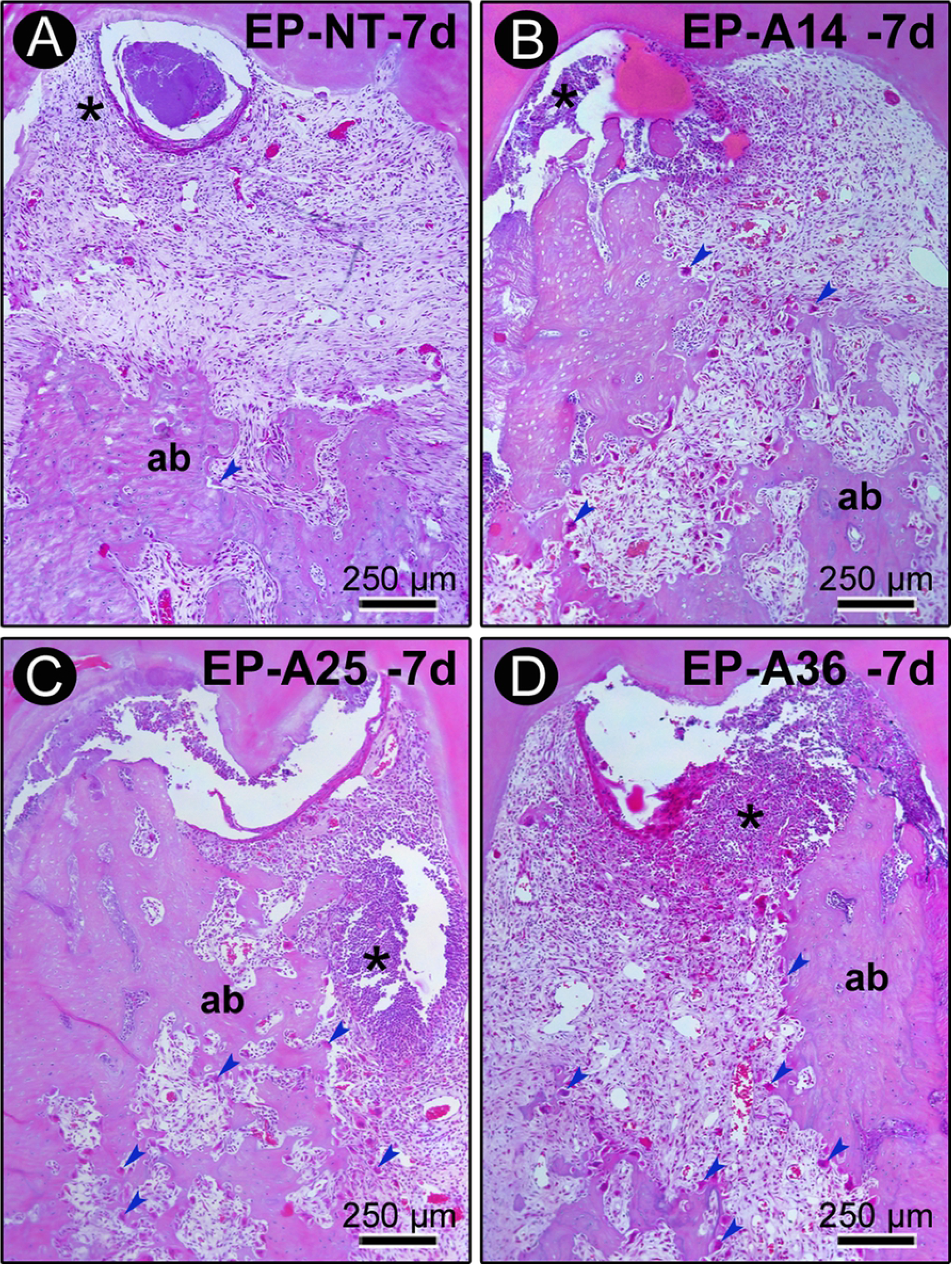
Histopathological characteristics of the furcation region of the mandibular first molars of rats with PE in groups EP (A), EP-A14 (B), EP-A25 (C) and EP-A36 (D) 7 days after ligature placement. Photomicrographs showing the inflammatory infiltrate and alveolar bone loss. Abbreviations and symbols: ab, alveolar bone; *, inflammatory infiltrate. Staining: hematoxylin and eosin (HE). Magnification: 100x. Scale bar: A – d: 250 μm.

At days 15 and 30 after ligature placement, chronic inflammatory infiltrate was present in the connective tissue of the furcation region of the EP-NT group, and the alveolar bone loss had stabilized. At day 15 in the groups treated with alcohol, the inflammatory response and bone resorption activity were still very intense. In groups EP-A14 and EP-A25, a mild inflammatory response was observed only after 30 days (Figure 4). During this period, these groups still had areas of active bone resorption. In the EP-A36 group, the local inflammatory response and bone resorption activity were still exacerbated at 30 days compared with the other groups.

**Figure 4:**
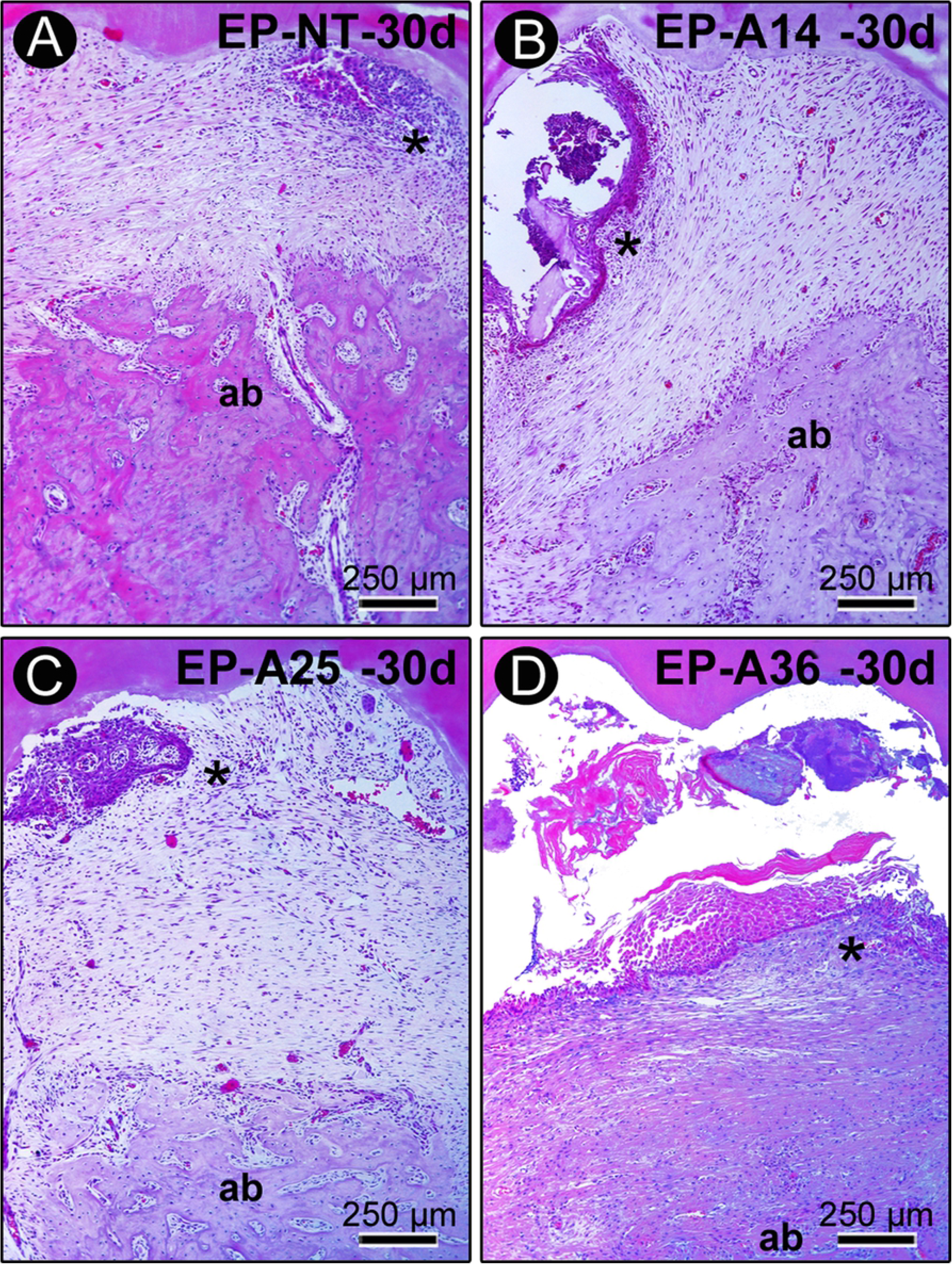
Histopathological characteristics of the furcation region of the mandibular first molars of rats with PE in groups EP (A), EP-A14 (B), EP-A25 (C) and EP-A36 (D) 30 days after ligature placement. Photomicrographs showing the inflammatory infiltrate and alveolar bone loss. Abbreviations and symbols: ab, alveolar bone; *, inflammatory infiltrate. Staining: hematoxylin and eosin (HE). Magnification: 100x. Scale bar: A – d: 250 μm.

### 3.3. Histometric analysis

The Kappa test indicated a level of intraexaminer agreement for the PBF measurements of 94%, which was a high level of agreement.

The results are shown in Table 1. According to the intragroup analysis comparing the different periods, the EP-A14, EP-A25 and EP-A36 groups had lower PBFs at days 7, 15 and 30 compared with the values obtained at day 3 (P ≤ 0.05). According to the intergroup analysis comparing the different groups, the animals of groups EP-A14, EP-A25 and EP-A36 had lower PBFs than the animals belonging to the EP-NT group at days 7, 15 and 30 (P ≤ 0.05). At day 3, there were no significant differences between the groups, and there were no significant differences between the EP-A14, EP-A25 and EP-A36 groups at any time point.

**Table 1:** **A:** Means and standard deviations (M ± SD) of the histometric data for PBF (%) in the furcation regions of the left mandibular first molars according to groups and periods. **B:** Means and standard deviations (M ± SD) of the amount of TRAP in the furcation regions of the left mandibular first molars according to groups and periods. Symbols: *statistically significant difference with EP-NT group at the same point. ^†^ statistically significant difference with EP-A14 group at the same point. ^&^statistically significant difference with EP-A25 group at the same point. ANOVA and Tukey test (p≤0.05).

| A |  | 3 DAYS | 7 DAYS | 15 DAYS | 30 DAYS |
| --- | --- | --- | --- | --- | --- |
|  | EP-NT | 77.98±2.62 | 77.12±3.28 | 74.05±3.31 | 75.04±2.27 |
|  | EP-A14 | 77.54±2.27 | 57.45±12.24* | 48.62±22.44* | 51.51±13.31* |
|  | EP-A25 | 75.81±2.85 | 55.16±15.11* | 51.48±14.61* | 50.91±2093* |
|  | EP-A36 | 77.67±3.05 | 48.74±12.01* | 34.82±5.49* | 34.50±7.61* |
| B |  |  |  |  |  |
|  | EP-NT | 10.94±4.33 | 18.20±10.25 | 9.70±6.95 | 8.94±4.74 |
|  | EP-A14 | 30.88±7.48* | 31.60±3.78* | 34.58±9.12* | 38.98±5.33* |
|  | EP-A25 | 31.76±8.19* | 38.40±5.45*,† | 31.80±2.92* | 35.04±5.88* |
|  | EP-A36 | 57.64±5.21*,†,& | 50.60±4.60*,†,& | 54.47±7.62*,†,& | 59.00±4.73*,†,& |

### 3.4. Immunohistochemical analysis

The RANKL and OPG results are shown in Figure 5. For RANKL, the EP-NT group exhibited high immunoreactivity pattern at days 3 and 7 and moderate immunoreactivity pattern at days 15 and 30. The EP-A14 group exhibited high immunoreactivity pattern at days 3, 7 and 15 and moderate immunoreactivity pattern at day 30. The EP-A25 and EP-A36 groups exhibited high immunoreactivity pattern at all time points. The OPG immunoreactivity pattern was predominantly low at all time points in all the groups.

**Figure 5:**
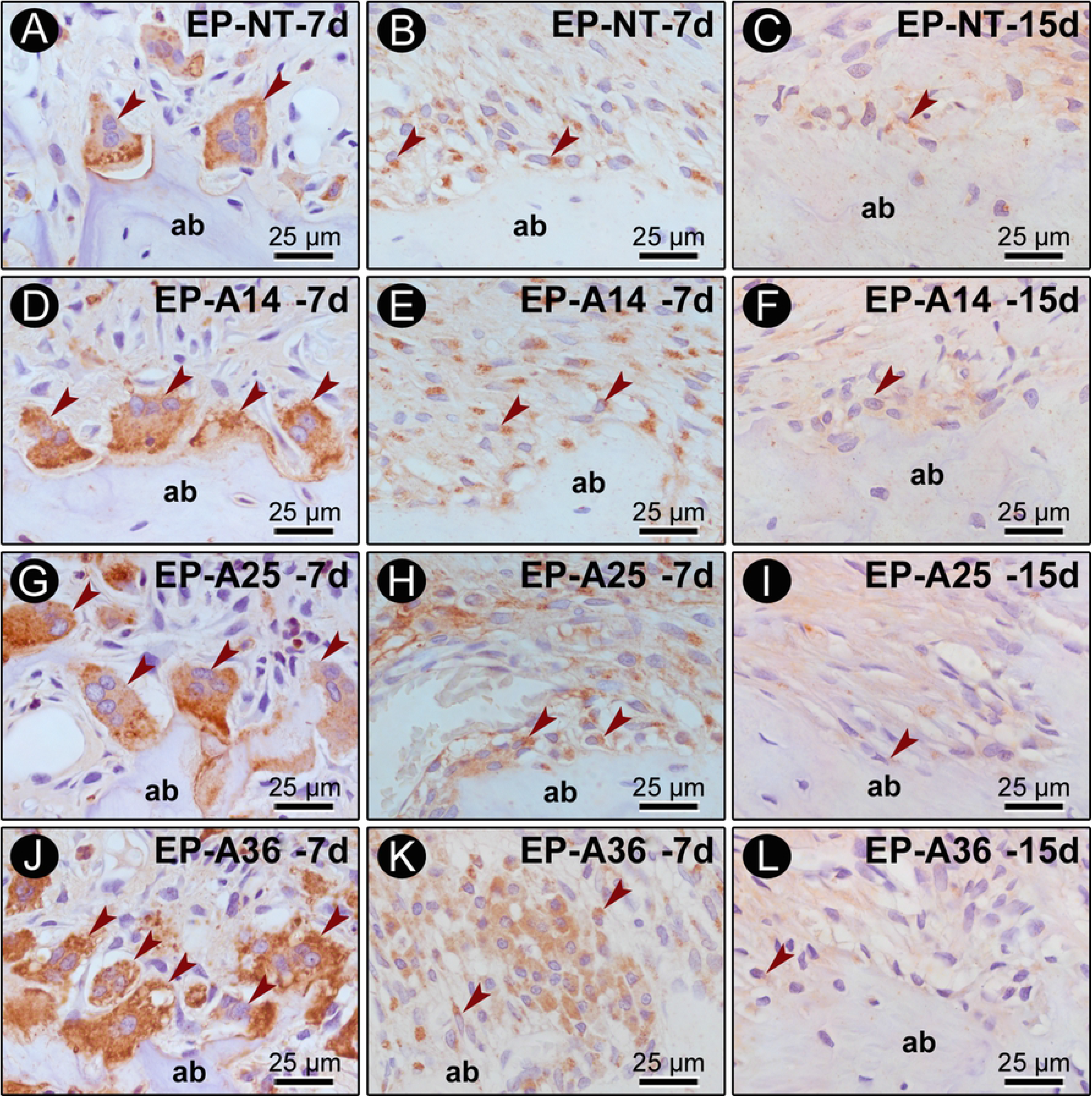
TRAP, RANKL and OPG immunostaining in the furcation region of the mandibular first molar. Photomicrographs showing the immunostaining pattern of TRAP (A, D, G, J)), RANKL (B, E, H, K)) 7 days after ligature placement, and OPG (C, F, I, L) 15 days after ligature placement. Abbreviations: ab, alveolar bone; arrows, immunostaining cells. Magnification: 1000x. Scale bars: 25 μm. Counterstaining: hematoxylin.

The TRAP results are shown in Table 1 and Figure 5. According to the intergroup analysis comparing the different experimental groups, the EP-A14, EP-A25 and EP-A36 groups had a higher number of TRAP cells than the EP-NT group at all time points (P ≤ 0.05). The EP-A25 group had a higher number of TRAP cells than the EP-A14 group at day 7, and the EP-A36 group had a higher number of TRAP cells than the EP-A14 and EP-A25 groups at all time points (P ≤ 0.05).

## 4. Discussion

The hypothesis that chronic alcohol consumption would increase the severity of EP regardless of the concentration consumed was confirmed in our study. This study demonstrate that chronic alcohol consumption increases the severity of EP in a dose-dependent manner by increasing the magnitude of the local inflammatory response, RANKL immunoreactivity pattern and the number of TRAP+ cells, which leads to greater alveolar bone loss and a lower PBF.

Various methodologies, which differed in regard to the way the alcohol was administered, the concentration of the alcohol solution, the age of the animals and the type of alcohol, have been applied to evaluate the effects of alcohol on EP [27,34–37]. Consequently, the lack of methodological standardization has given rise to conflicting results. For this reason, the animals in the present study were exposed to different alcohol concentrations (14%, 25% and 36%) to determine which concentration could demonstrate a positive association between alcohol consumption and periodontal disease.

The experimental model of chronic alcoholism used in this study was proposed by Souza et al., (2009) [34]. A 30-day period of prior exposure to alcohol in rats mimics the behavior of a chronic alcoholic [34] and has been proven to be an effective experimental model of alcoholism [38,39]. This model is easy to implement and extremely practical, especially when long experimental periods are required. It also involves virtually no manipulation of the animal, as is often required in other experimental models. In this study, the serum ALT and AST levels confirmed the liver damage caused by chronic alcohol consumption in the groups exposed to alcohol, as changes in liver enzymes have been observed in alcoholic patients [40].

The effects of alcohol on immune responses depend on the timing of consumption. In acute consumption, the anti-inflammatory effects are due to a reduction in pro-inflammatory cytokines and an increase in anti-inflammatory cytokines, which suggests a protective effect [41]. In chronic use, alcohol acts in an opposite way by increasing the levels of pro-inflammatory cytokines, such as tumor necrosis factor (TNF), interleukin 1 (IL-1) and interleukin 6 (IL-6), and reducing the levels of anti-inflammatory cytokines [17,42]. In our results, the inflammatory response in PE was more exacerbated in alcohol dose-dependent manner, corroborating these studies.

The results of the present study revealed that in the EP-NT group, comprising animals that were not exposed to alcohol, there was a gradual increase in inflammatory infiltrate 3 and 7 days after ligature placement and a subsequent reduction in its magnitude as the inflammatory process progressed as well as a concomitant reduction in the PBF. These results agree with previous studies [43,44]. Similarly, the EP-A14, EP-A25 and EP-A36 groups, which consisted of animals that were exposed to alcohol, exhibited similar histopathological characteristics. However, the magnitude of the local inflammatory responses was much higher, especially in the groups that were exposed to higher alcohol concentrations, and these groups also had large areas of bone necrosis.

There was a gradual reduction in the PBF over time in all the groups that were exposed to alcohol. However, there were no significant differences among the groups of animals that were exposed to alcohol. Such histological and histometric findings demonstrate that chronic alcohol use greatly exacerbates the severity of EP, irrespective of the concentration was used. In contrast to our results, a study by Liberman et al., (2011) [27] revealed that alcohol intake did not affect alveolar bone loss in EP. However, the concentration used in their study (5%) is considered low and was below the lowest concentration used in our study (14%). Our data corroborate previous studies that used concentrations ranging from 10% to 30% and found that chronic alcohol consumption caused alveolar bone loss and decreased bone density in animals with EP [35,45,46].

Periodontal bone remodeling is locally regulated by the RANK-RANKL-OPG system, which modulates alveolar bone resorption. RANK, its ligand RANKL and OPG are included in this system [47]. Thus, the resorptive activity of alveolar bone in this study was evaluated by using immunohistochemistry to detect the 2 key regulators of local bone metabolism, RANKL and OPG, and by quantifying the TRAP+ multinucleated cells. Previous experimental studies have evaluated the effects of alcohol consumption on bone metabolism and osteoclastogenesis and noted a decrease in osteoblast activity [48] and an increase in osteoclastic activity [49]. In the present study, the OPG immunostaining was consistently low in all the experimental groups and at all the time points. In contrast, RANKL immunoreactivity pattern showed a dose-dependent increase when compared to the EP-NT group, especially 3 and 7 days after ligature placement. Corroborating the OPG and RANKL results, we observed that the groups exposed to chronic alcohol had higher numbers of TRAP+ multinucleated cells in the furcation region than the EP-NT group. The EP-A36 group had a higher number of TRAP+ cells than the EP-A14 and EP-A25 groups. The same result was found in a recent study that evaluated apical periodontitis in animals with chronic alcohol consumption at a concentration of 20° [22]. These results therefore suggest that the regulation of osteoclastogenesis in animals subjected to chronic alcohol consumption is dose dependent.

The way in which chronic alcohol consumption stimulates osteoclastogenesis has been studied by Iitsuka et al., (2012) [48]. They concluded that alcohol stimulates osteoclastogenesis by increasing RANK expression, which is mediated by the production of reactive oxygen species and the activation of extracellular signal-regulated kinases in precursor osteoclast cells and results in an increase in RANKL in osteoblasts. In the pathology of periodontal disease, a series of cascading events leads to osteoclastogenesis, which is regulated by the RANK-RANKL-OPG system [32,47]. We can therefore infer that chronic alcohol consumption can worsen the severity of periodontal disease. In this context, the results of our study demonstrate that chronic alcohol consumption can worsen alveolar bone loss in EP via RANKL upregulation. In contrast to our results, Bastos observed a significantly lower number of TRAP+ cells, as well as an increase in OPG, in a group with induced EP that was exposed to alcohol compared with the control group [34] (Bastos et al., 2011). However, an increase in RANKL was observed in the same animals with EP that were exposed to chronic alcohol consumption when compared with the control group. It is important to note that in the same study [35], increased bone loss and decreased bone density were observed in animals with EP that were exposed to chronic alcohol consumption, confirming the deleterious effects of chronic alcohol consumption on the resorptive activity of alveolar bone.

The histological and immunohistochemical findings of our study indicate that chronic alcohol consumption has dose-dependent effects. In the animal groups that were exposed to alcohol, higher concentrations resulted in more severe inflammation, increased RANKL immunoreactivity and a significantly higher number of TRAP+ cells. However, the magnitude of the immunoinflammatory responses did not result in statistically significant changes in the PBF among the animal groups that were exposed to alcohol, although it may have had significant effects on bone tissue in later periods. It is important to note that pre-clinical studies using animals are necessary to protect human health and guide clinical studies investigating therapeutic strategies for the treatment of periodontal disease.

## 5. Conclusion

Within the limits of the present study, we can conclude that chronic alcohol consumption at different concentrations increases the severity of EP regardless of the concentration consumed in a dose-dependent manner by increasing the magnitude of the local inflammatory response and stimulating alveolar bone resorption.

## Compliance with ethical standards

### Funding

We thank the Brazilian Federal Agency for the Support and Evaluation of Graduate Education (Coordenação de Aperfeiçoamento de Pessoal de Nível Superior - CAPES) for the scholarship and the Fundunesp-PROPE 0096/004/13 for assistance in the development of this research.

### Ethical approval

All applicable international, national, and/or institutional guidelines for the care and use of animals were followed. All procedures performed in studies involving animals were in accordance with the ethical standards of the institution or practice at which the studies were conducted.

### Informed consent

For this type of study, formal consent is not required.

